# Are all vet schools equal? An exploration of postgraduate qualification attainment by alumni cohorts of seven UK veterinary schools

**DOI:** 10.64898/2026.01.03.697366

**Authors:** Peers Davies, Alexander Corbishley, Joseph Neary

**Affiliations:** Department of Livestock & One Health, Leahurst Campus, University of Liverpool, CH64 7TE; The Royal (Dick) School of Veterinary Studies and The Roslin Institute, Easter Bush Campus Midlothian EH25 9RG

## Abstract

Do graduates from the different vet schools attain clinical and academic postgraduate qualifications (certificates, diplomas, masters, PhD and Fellowship) at the same rates? This is important because leadership and progress within the veterinary profession, as in human medicine, comes from advancing the frontier of our knowledge through research and clinical specialisation. Postgraduate qualifications are essential training for both and are therefore useful metrics to measure. In this study the Royal College of Veterinary Surgeons annual registers of qualified vets from 2000-2021 were analysed. Significant and substantial differences in the proportion of graduates from different universities attaining postgraduate qualifications were observed. Whilst associations identified by this analysis cannot prove causation, they do strongly suggest the wide range of university-specific factors, such as student selection criteria, teaching methods, curriculum design and assessments which contribute to the culture and ethos of the institution have an impact on the career trajectory of their graduates.

## Methodology

Ethical approval was not required for analysis of open access public datasets. An electronic copy of the Royal College of Veterinary Surgeons (RCVS) register [1] was provided by the RCVS for 2000 – 2021 (except 2002 due to being unavailable in an electronic format). The register contained individual records for each member and fellow of the RCVS including their undergraduate veterinary qualifications and the university from which it was awarded and the year in which they first registered with the RCVS (which is not necessarily synonymous with the year of graduation). The register also contains a list of all postgraduate qualifications attained and declared voluntarily by each member, but not the institution where, or date when, they were awarded. Data were organised in Microsoft Excel and statistical analyses were performed in Minitab22 [2] and R-4.5.2 [3]. The type and number of postgraduate qualifications per person were calculated, excluding bachelors or primary veterinary qualifications including the VetMB required for registration or honorary qualifications such as MA(Cantab) or HonFRCVS to only recognise earned qualifications. FRCVS status was modelled as a separate outcome variable from other postgraduate qualifications because of the diverse range of subjective criteria of which it can be awarded.

MRCVS and FRCVS vets who qualified from non-UK universities and the University of Surrey (first cohort only graduated in 2019) graduates were removed. This left 22,570 vets graduating from the universities of Bristol, Cambridge, Edinburgh, Glasgow, Liverpool, London (RVC) and Nottingham on the 2021 register. Date of first registration ranged from 1944 to 2021. The percentage of alumni with a postgraduate academic or specialist clinical qualification was calculated per university along with the average number of postgraduate qualifications per vet per university. To assess the retention of graduates in the profession post-graduation, the number of first registrations in each of the available years from 2000 to 2020 was used as a proxy for the number of graduates per institution per year and as a total for the period. This was compared to the number of graduates still registered in 2021 who first registered in each year of the prior two decades. This crude proxy does not account for first registrations in a subsequent year to that in which a person graduated. This may happen with graduates moving abroad to practice and subsequently returning. To address this limitation a second analysis tracked each individual who first registered in 2011 (proxy for year of graduation) through the subsequent 2016 and 2021 registers to calculate retention on the register and retention as ‘UK Practicing’ vets on the register at five years and 10 yrs post first registration.

To account for the bias associated with the number of years since graduation on the likelihood of attaining a postgraduate qualification, a multi-variable analysis was used to account for the years since first registration. This also accounted for the differing age demographics of the alumni populations of the different universities, which has come about through the differing expansion patterns of student populations between institutions. The relative number of alumni from each university with a postgraduate qualification was analysed in R using a series of binary logistic regression models with the award of FRCVS, MSc and PhD as individual outcomes, and combined outcomes for clinical certificates or diplomas (clinical specialist qualification) and combined MSc & PhD (academic qualifications) as outcome variables. University was coded as a categorical predictor and number of years since first registration as a continuous predictor. The register did not contain sex/gender information so this could not be included. The registration category (UK practicing, Practicing outside the UK and Non practicing) was assessed as a categorical predictor, but not included in the final model, as the impact on other estimates and model fit were negligible. Model fit was assessed by ROC/AUC, AIC and Hosmer-Lemeshow test. The number of postgraduate qualifications per graduate was analysed as a zero inflated negative binomial model using the ‘pscl’ R package [4].

## Results

The retention rate of graduates from the register was largely consistent across the UK universities and over time, averaging 95%, with a larger apparent drop in the number of retained graduates from the Scottish Universities (89% Edinburgh, 81% Glasgow), which can be accounted for by their higher number of overseas students who may only have worked in the UK temporarily. Three universities recorded apparent positive retention rates of 101%-105% indicating that the assumption that graduation year and first registration year are synonymous leads to misattribution of graduates to graduation years in a low but non-trivial percentage of cases as described in Methods. In the five- and ten-year post first registration analysis using the 2011, 2016 and 2021 data following individual registration identifiers, the retention rate was again similar between the English universities and lower for Edinburgh and Glasgow both in all registration categories.

The proportion of recent graduates (first registered 2011-21) per university awarded one or more postgraduate qualifications by 2021 averaged 10.1%, with a wide variation between institutions from 7.9% of Glasgow graduates to 13.5% of Cambridge graduates. The denominator for this calculation being those graduates who were on the 2021 register, therefore removing any bias associated with some universities accepting higher proportions of international students who practice abroad after graduating and do not remain on the RCVS register long term. Among all graduates (first registered 1944-2021) appearing on the 2021 register, the proportion with one or more postgraduate qualification increased to an average of 24.7% across all universities and the disparity between institutions grew substantially, with 34.5% of Cambridge graduates achieving at least one postgraduate qualification, significantly higher than any of the other schools (range 22%-26%; p<0.01). A detailed breakdown, including the average number of postgraduate qualifications per person by type of qualification and cohort is provided in Table1.

After accounting for the age demographic variation between institutions, the binomial logistic regression model highlighted a substantially and significantly higher odds of Cambridge alumni acquiring both specialist clinical and academic postgraduate qualifications, particularly PhDs (Figure 1), with ROC/AUC of 0.63 & 0.74 respectively. Graduates from Bristol, Edinburgh, Glasgow, Liverpool and London attained at similar rates, significantly ahead of Nottingham alumni. The number of postgraduate qualifications per graduate, accounting for graduate age did not vary significantly between institutions as evaluated in the zero inflated negative binomial model. The proportion of graduates awarded fellowship of the RCVS (FRCVS) was substantially lower than for postgraduate qualifications at 1.7% (374 of 22,570) among the universities Cambridge and Bristol had a significantly higher proportion of FRCVS graduates at 2.9% and 2.2% respectively.

**Figure 1.**
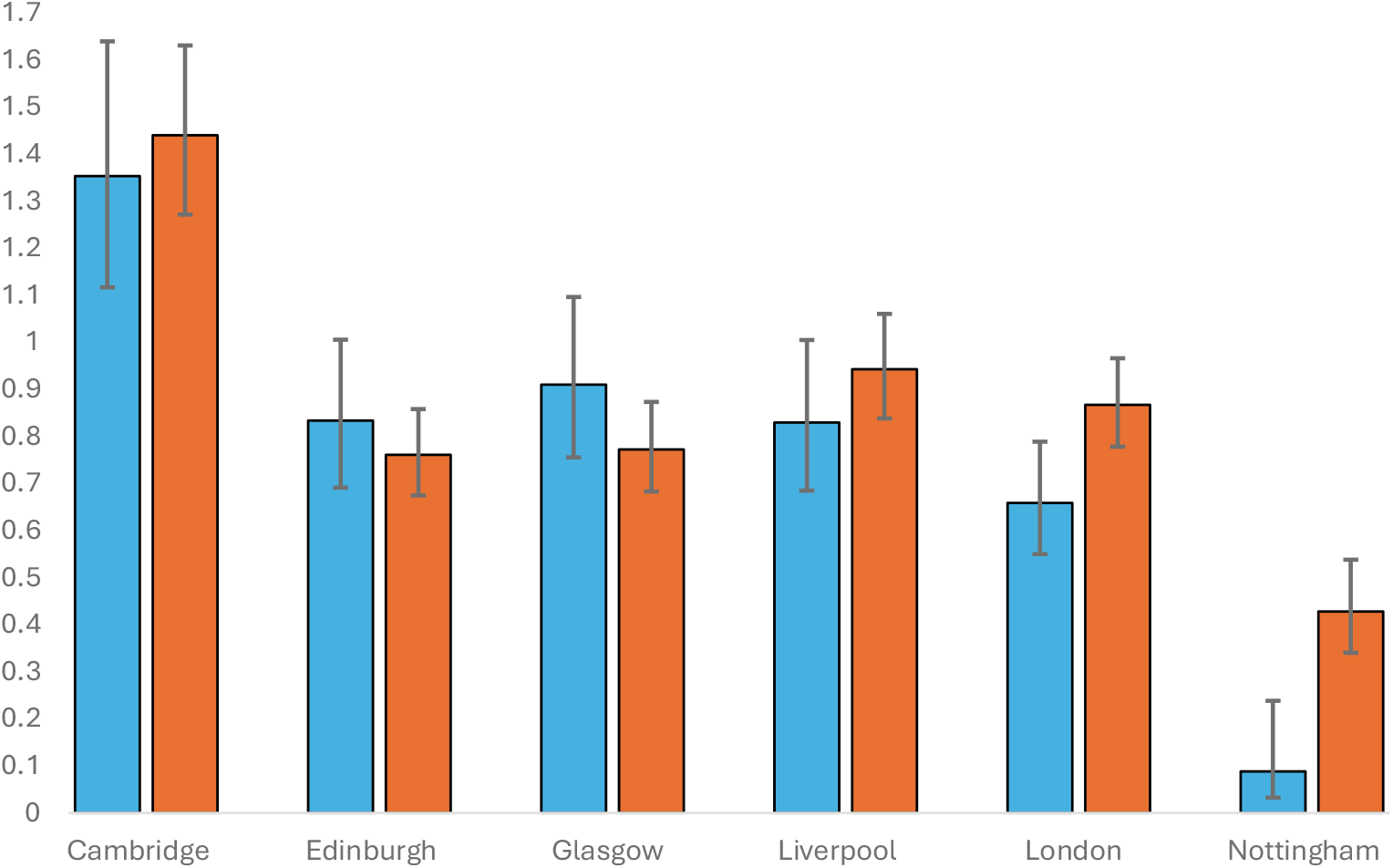
Odds ratios and 95% confidence limits for separate binomial logistic regression models of the award of one or more postgraduate academic qualifications (model 1: MSc & PhD, BLUE) and/or one or more postgraduate clinical specialist qualifications (model 2: Certificate and Diploma, ORANGE) to alumni who received their undergraduate veterinary degree from each UK veterinary school relative to University of Bristol as the reference category i.e. OR=1.0. NB. The data relates to all members of the 2021 register with a ‘years registered’ independent variable from 1944 to 192. However, the first Nottingham cohort graduated 2011, therefore their results relative to the other universities should be interpreted with caution and considered alongside the results in Table 1.

**Table 1.** Comparison between Universities of the average number of postgraduate qualifications (MSc, PhD, Certificate & Diploma) per graduate for all graduates on the 2021 RCVS register and for recent graduates defined as those first appearing on the register on or after 2011; the percentage of graduates with one or more academic postgrad qualifications (MSc or PhD) and the percentage of graduates with one or more specialist clinical postgrad qualifications (Certificate or Diploma). The recent graduate population represents the most comparable dataset for Nottingham graduates.

|  | All Graduates (1944 – 2021) |  |  | Recent Graduates (2011-2021) |  |  |
| --- | --- | --- | --- | --- | --- | --- |
|  | mean<br>postgrad<br>quals per vet | % of vets<br>with a MSc<br>or PhD | % of vets<br>with a Cert<br>or Dip | mean<br>postgrad<br>quals per vet | % of vets<br>with a MSc<br>or PhD | % of vets<br>with a Cert or<br>Dip |
| Bristol | 0.39 | 7.5% | 21.9% | 0.14 | 1.8% | 10.1% |
| Cambridge | 0.54 | 10.5% | 29.5% | 0.16 | 1.4% | 12.6% |
| Edinburgh | 0.32 | 7.0% | 18.2% | 0.09 | 1.4% | 7.0% |
| Glasgow | 0.35 | 8.3% | 19.1% | 0.09 | 0.9% | 7.1% |
| Liverpool | 0.35 | 6.5% | 21.0% | 0.14 | 1.6% | 10.6% |
| London | 0.33 | 5.3% | 19.7% | 0.12 | 1.1% | 9.1% |
| Nottingham | 0.10 | 0.4% | 8.6% | 0.10 | 0.4% | 8.6% |

## Discussion

The RCVS register data provides a useful insight into aspects of post-graduate career progression. The main focus of this study was to explore the association between the vet school attended and the likelihood of attaining higher academic or specialist clinical qualifications. Obtaining postgraduate qualifications has never been a mandatory requirement in a veterinary career, and many vets have historically enjoyed very successful and fulfilling careers without further study. However, it is hard to make a case that attaining postgraduate qualifications is anything other than a good thing for both the individual and the profession as a whole. The results of this analysis demonstrate that postgraduate attainment is associated with the university attended as an undergraduate. Specifically, Cambridge graduates appear more likely to pursue further study than graduates of the other universities. The data available does not allow any determination of causality, although it would be reasonable to speculate that student entry selection criteria, curriculum and teaching methods may all play a part. Cambridge has long emphasised a particularly rigorous and demanding scientific education, equipping veterinary students with a firm grasp of the fundamental underpinning biology of the species and diseases they will treat as qualified vets. The additional compulsory year of intercalated study in a discipline of the student’s choice provides an immersive experience of conducting research and research led teaching, which is only pursued by a very small minority of students in other universities who must deliberately opt for intercalation. All these factors may contribute to the subsequent higher postgraduate attainment of Cambridge graduates. The data contained within the register can only provide two predictor variables, which limits model fit and clearly cannot provide a comprehensive understanding of all the various factors and confounding variables which facilitate, encourage or deter veterinary graduates from pursuing specialist clinical, scientific and research qualifications. However, identifying real differences in university graduate career progression is useful for universities to reflect upon their course design, learning and development objectives, teaching methods and student selection criteria. It is also useful for the RCVS as the regulatory body responsible for accreditation of courses to appreciate that different teaching and selection methods can lead to different and better outcomes. This is especially true as the RCVS seeks to actively promote further postgraduate study and continual professional development. The results of this study are consistent with the high academic entry standards and research-focused education of Cambridge veterinary graduates.

In December 2025 the University of Cambridge announced they are considering closing their veterinary school. If this happens, what might be lost? Despite being smaller than the other schools, its graduates are over-represented at the cutting edge of scholarship and research and the veterinary school has been consistently ranked highly in national and international league tables. This makes it all the more disappointing that the veterinary school is threatened with closure when by this, and many other metrics, it is one of the best veterinary courses in the world.

## Acknowledgements

The author would like to thank the RCVS library team for providing the data.

## Data availability

All data and code used for analysis is available from the author upon request or from the University of Liverpool repository in due course following peer reviewed publication.

## Research ethics & funding statement

Ethical approval was not required for this analysis of open access public datasets. No funding was received for this project.

## Author Contributions

PD designed and conducted the data analysis and prepared the manuscript. AC edited the manuscript and AC & JN contributed to the analysis and interpretation.

